# Long-term prevention of aneuploidy in human pluripotent stem cells by fine-tuning GSK3 activity

**DOI:** 10.1101/2025.10.27.684724

**Authors:** A. De Jaime-Soguero, M. Romitti, S. Costagliola, A. Jauch, G. Vilangappurath, K. Willert, F. Foijer, S.P. Acebrón

## Abstract

Human pluripotent stem cells (hPSCs) are critical cell sources to model human development and hold promise towards regenerative medicine. However, cultured hPSCs quickly acquire chromosome gains or losses (aneuploidy) due to high replicative stress and errors in chromosome segregation, which hampers their use for stem cell therapies. We have recently described that embryo patterning signals control chromosome segregation fidelity during early human lineage specification by modulating the response to replicative stress. Here, we demonstrate that fine-tuned activation of WNT signalling using off-the-shelf GSK3 inhibitors prevents chromosome missegregation and aneuploidy during long term passaging of hPSCs. Lineage-specification and sequencing analyses demonstrate that hPSCs culture with E8 supplemented with a low dosage of GSK3 inhibitors retain primed pluripotency and full potential to the differentiation into the 3 germ layers for over 30 passages. Strikingly, detailed mitotic studies revealed that long-term cultured hPSCs often display ultra-fine chromosome bridges during anaphase due to unreplicated DNA, which can result in hitherto uncharacterised copy-number-variations and other genomic aberrations. We show that culture with low dosage of the GSK3 inhibitor CHIR99021, but not direct attenuation of DNA replication stress using nucleosides, prevents mitotic DNA synthesis and the formation of ultra-fine bridges during long term hPSC passage. Taken together, we propose to enrich E8 culture media with 100 nM CHIR99021 to maintain euploidy in primed hPSCs, and to routinely map for genomic alterations, in addition to aneuploidy, for modelling and regenerative medicine studies.

## Introduction

Chromosomal mosaicism refers to the presence of karyotypically distinct cell populations: euploid and aneuploid cells. Aneuploidy is often associated with decreased cellular fitness, and it is rarely found in adult human tissues, with the exception of cancer [1, 2]. Strikingly, around 80% of human embryos in the first two weeks of development present chromosomal mosaicism [3-5], which is the main cause of miscarriage in humans [6, 7]. Deep sequencing studies indicate that early embryos (prior gastrulation) are also hotspots for genome mosaicism, most notably SNVs and CNVs [8-10]. Recent lines of evidence suggest that a dynamic DNA replication programme during early development is responsible for both structural and numerical chromosome aberrations in human early embryos [8, 11-16].

Chromosomal mosaicism is also a pervasive problem in 2D and 3D *in vitro* models for human early development. In particular, human pluripotent stem cells (hPSCs) often acquire chromosome aberrations over time, with the incidence rapidly increasing with passage number [17, 18]. Critically, gains in some chromosomes (1, 8, 12, 17 and 20) [18, 19], as well as recurrent deletions of specific chromosome arms (10p, 18q and 22q) [17, 18], provide growth advantage to hPSCs and often take over the cell population, which precludes their expansion in bulk, and instead requires lengthy and careful evaluation of single clones towards modelling studies and regenerative strategies [20]. In particular, frequent aneuploidies in PSCs are linked with alterations in their differentiation capacity towards the three germ layers [18, 21-23].

We have recently identified that basal DNA replication stress during pluripotency triggers i) increased microtubule polymerisation rates during the subsequent mitosis, leading to kinetochore-microtubule attachment errors and whole-chromosome missegregation; and ii) unreplicated and damaged DNA, resulting in ultra-fine DNA bridges during anaphase [24]. In addition, DNA replication stress causes chromatin condensation defects and subsequent aneuploidy during pluripotency [24, 25]. Together, these results explain the low chromosome segregation fidelity in human pluripotent stem cells [24, 26]. Previous studies proposed to supplement hPSC cultures with nucleosides to alleviate DNA replication stress and subsequent chromosome missegregation [24, 27], possibly by replenishing endogenous nucleotide pools after the firing of too many origins [27-30]. However, recent studies in mouse embryonic stem cells (mESCs) indicated that although prevent nucleosides the activation of replicative stress responses, do not ensure the timely completion of DNA replication prior mitosis [31].

Here, we demonstrate that fine-tuned activation of WNT signalling using off-the-shelf GSK3 inhibitors prevents chromosome missegregation and aneuploidy during long term passaging of hPSCs. Lineage-specification and sequencing analyses demonstrate that hPSCs culture with a low dosage of the GSK3 inhibitor CHIR99021 retain prime pluripotency and full potential to the differentiation for over 30 passages. Importantly, detailed chromosome segregation studies revealed that long-term culture of hPSCs often results in ultra-fine chromosome bridges due to unreplicated DNA, which can result in hitherto uncharacterised copy-number-variations and other genomic aberrations. We show that culture with 100 nM CHIR99021, but not with nucleosides, promotes the completion of DNA replication before G2/M, and prevents the formation of ultra-fine chromosome bridges during long term hPSC expansion. Taken together, we propose to reformulate hPSC culture conditions by including 100 nM CHIR99021 in E8 media, and to routinely map for structural chromosomal alterations, in addition to aneuploidy, for modelling and regenerative medicine studies.

## Results

We have recently identified that patterning signals regulating anteroposterior patterning during gastrulation also form part of a signalling rheostat modulating DNA replication stress and chromosome segregation fidelity during naive and primed pluripotency [24]. On the one hand, we found that FGF signals, which are critical for self-renewal and to maintain primed pluripotency, induce replication stress and subsequent chromosome missegregation in human pluripotent stem cells (hPSCs) [24]. On the other hand, we identified that the posteriorising signal WNT protect differentiating hPSCs from different sources of DNA replication stress [24], functioning downstream of stalled forks in S-phase [24, 32]. The standard formulation of the hPSC culture media E8 contains FGF2, among another factors. We hypothesised that modulation of the signalling rheostat regulating DNA replication stress, in particular uncoupling WNT roles in lineage specification and genome maintenance, could prevent chromosome missegregation in hPSCs, without triggering differentiation, thereby allowing for their safe long-term expansion.

The GSK3 inhibitor CHIR99021 is an off-the-shelf reagent often used in stem cell culture and human lineage specification experiments to robustly activate WNT signalling [24, 33-36]. In agreement of the role of WNT in the primitive streak induction [37], titration of CHIR99021 revealed a dose dependent effect on hiPSC pluripotency exit (Figure 1A). In particular hiPSCs cultured with 500 nM CHIR99021 rapidly differentiated, while hiPSCs supplemented with a low dosage of CHIR99021 (10 – 100 nM) still formed homogenous colonies after 5 passages in E8 media, similarly as cells treated with nucleosides (Figure 1A). Similarly, culture of hiPSCs for 30 passages in E8 supplemented with 100 nM CHIR99021 did not significantly impact the expression of the pluripotency gene *NANOG*, nor induced the expression of the primitive streak marker *T* (Brachyury) (Figure 1B).

**Figure 1:**
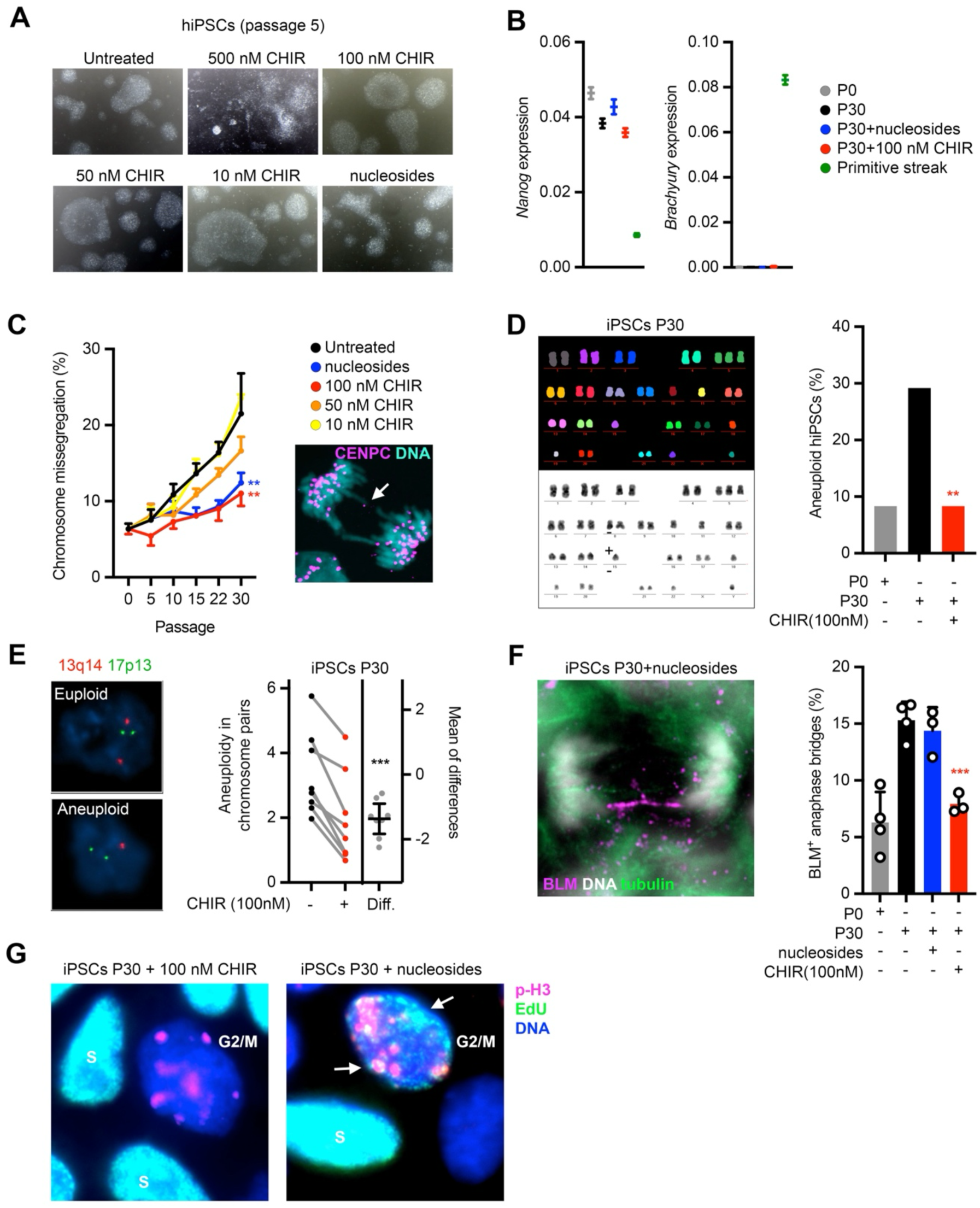
Chemical regulation of GSK3 activity prevents mitotic aberrations and aneuploidy in hiPSCs. **A**, Colony formation analyses of hiPSCs culture for 5 passages in E8 supplemented with the indicated factors. **B**, qPCR analyses of hiPSCs at passage 0 (P0) or 30 (P30), as well as in vitro generated human primitive-streak-like cells. **C**, Chromosome segregation studies of hiPSCs expanded in E8 with the supplemented with the indicated factors, and analysed at the indicated passages. **D**,**E**, M-FISH and interphase FISH studies of hiPSCs cultured as indicated **F**,**G**, Immunofluorescence studies in passage 0 or 30 hiPSCs cultured in E8 supplemented with the indicated factors. CHIR99021 (CHIR).

We culture of hiPSCs for 30 passages with different concentrations of CHIR99021 or in presence of nucleosides, and analyse their ability to properly segregate chromosomes in mitosis by focusing on the presence of lagging chromosomes during anaphase (Figure 1C). Untreated hiPSCs decrease chromosome segregation fidelity with each passage, reaching a striking 23% of anaphases with lagging chromosomes at passage 30 (Figure 1C). Importantly, supplementation with 100 nM CHIR99021 or nucleosides during long-term expansion largely protect hiPSCs from the increase in chromosome missegregation (Figure 1C). Of note, we observed similar results in other hiPSC lines, as well as in H1 and H9 hESCs (Figure S1A).

KaryoFish analyses of metaphase spreads revealed that early hiPSC passages display 8% aneuploid cells, similarly to previous estimates [19, 24, 25], which increased up to 29% of hiPSCs at passage 30 (Figure 1D). Critically, hiPSCs maintained with 100 nM CHIR99021 did not show any increase in aneuploidy rates compared to early passages (Figure 1D). These results were backed up by interphase FISH analyses showing an average decrease in aneuploid rates of 1.4% per each investigated chromosome pair in the 100 nM CHIR99021 expanded hiPSCs (Figure 1E and S1B).

DNA replication stress can induce both numerical and structural chromosome instability [24, 29, 32, 38]. Structural and partial chromosome errors can often be tracked to DNA replication stress arisen from slowed or stalled replication forks during S-phase, as well as to impaired DNA repair, which can lead to ultra-fine bridges during anaphase [24, 38-40]. Strikingly, long-term passaged hiPSCs suffered from an elevated proportion of ultra-fine DNA bridges during anaphase (Figure 1F). Of note, expansion of hiPSCs during 30 passages with 100 nM CHIR99021, but not with nucleosides, prevented this structural chromosomal aberration (Figure 1F). These results unravelled a hitherto uncharacterised mitotic aberration in long-term culture hiPSCs which can be addressed by partial GSK3 inhibition, possibly due to WNT signalling roles managing different sources of DNA replication stress [24], instead of only counteracting nucleotide pool deprivation, as in the case of nucleosides [27, 28, 31].

Of note, RNA sequencing experiments showed no meaningful differences in cell fate between early and late hiPSC passages, including cells expanded with 100 nM CHIR99021, specially in comparison with primitive streak-like cells (Figure 2A-C). In agreement with a post-translational role of WNT signalling in genome stability [24, 32, 41, 42], hiPSCs passaged with or without 100 nM CHIR99021 barely display any differentially expressed genes (Figure S2A). Intriguingly, among those few differentially expressed genes emerged APOBEC factors (APOBEC3G, APOBEC3F), which have been implicated in replicative stress management in cancer cells and could mediate this response [43, 44].

**Figure 2:**
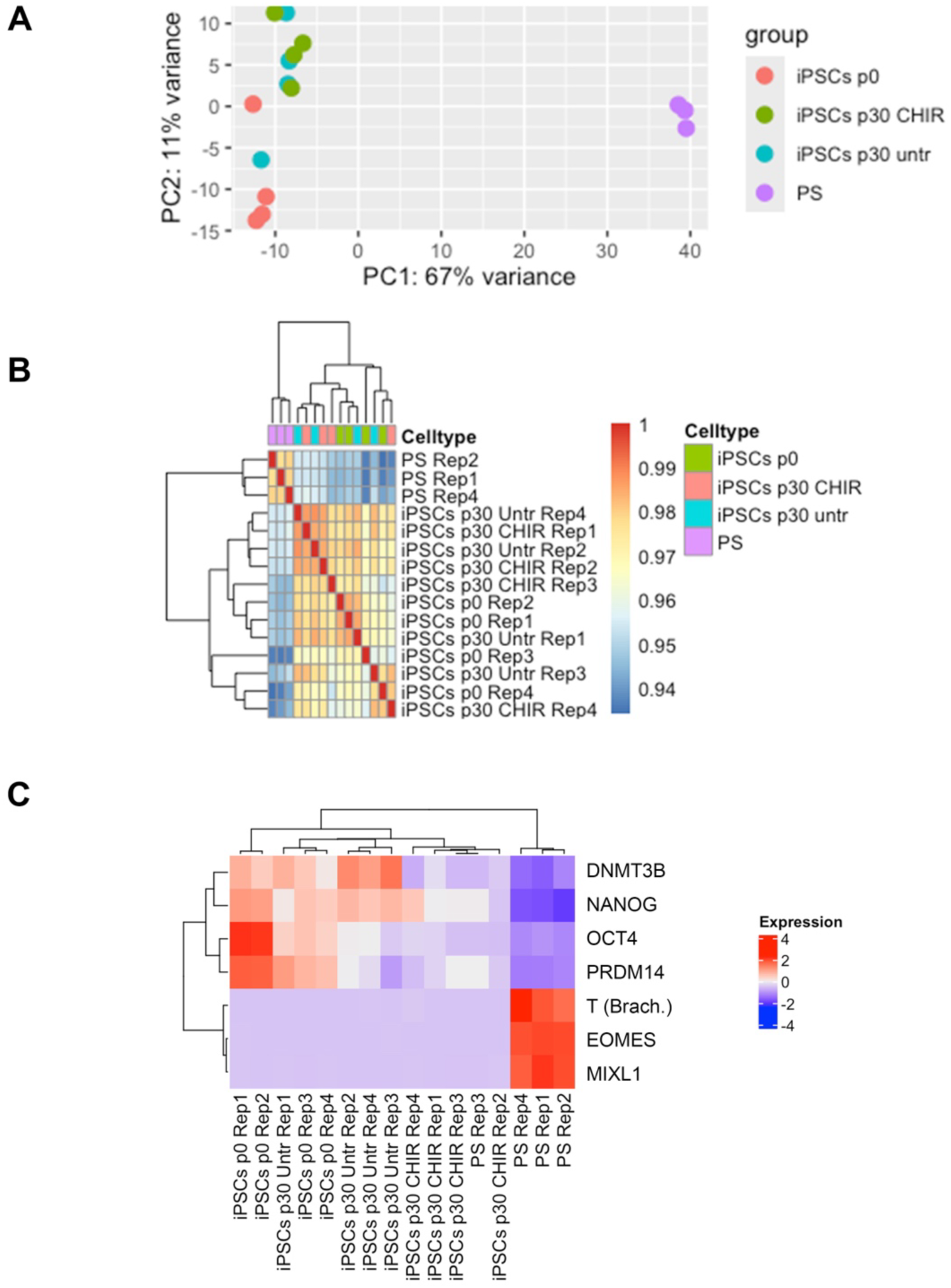
100 nM CHIR99021 does not impact hiPSC cell fate. **A-C**, Bulk sequencing analyses of human primitive-streak-like cells (PS), and hiPSCs cultured for up to 30 passages in E8 in the presence or absence of 100 nM CHIR99021 (CHIR).

Taken together, these results demonstrate that hPSC culture in E8 media supplemented with 100 nM CHIR99021 prevents whole-chromosome missegregation and unreplicated DNA during mitosis, thereby promoting numerical and structural chromosomal maintenance during long-term expansion of hPSCs.

## Discussion

The prevalence of chromosomal mosaicism in hPSCs is a major roadblock for their use in cell therapies, by requiring careful clone selection and karyotyping, instead of expansion in bulk. Previous research identified DNA replication stress as a critical culprit of chromosome segregation errors in in early embryos [8, 11-16] and hPSCs [24, 25, 27], including during reprogramming of iPSCs [30]. Attempts to alleviate this issue include overexpression of CHK1, as well as supplementation with exogenous nucleosides [27, 30]. Here, we demonstrate that mild activation of WNT signalling using off-the-shelf GSK3 inhibitors prevents chromosome missegregation and aneuploidy during long term passaging of hPSCs, without triggering differentiation or affecting their pluripotency potential. Critically, we also identified that long-term expansion of hPSCs results in other replication stress-associated phenotypes, including unreplicated DNA in G2/M, and subsequent ultra-fine chromosome bridges during anaphase. We showed that chemically fine-tuning GSK3 activity, but not exogenous nucleoside supplementation, ensures the completion of DNA replication before G2/M, thereby protecting hPSCs from other genomic aberrations. The lack of effect by nucleoside supplementation could be explained by recent evidence from mESCs, where they cause a miscoordination between the completion of DNA replication and cell cycle progression [31].

Our results highlight the opportunity for pharmacological modulation of patterning signalling pathways to ensure genome stability for stem cell and lineage specification models. In particular, off-the-shelf GSK3 inhibitors arise as a simple strategy to prevent chromosome and genome instability in hPSCs, opening the possibility of their safe long-term expansion. Of note both abnormally elevated and low WNT activity are linked with chromosomal instability [45-47], suggesting that WNT/GSK3 signalling adheres to a “Goldilocks principle”, in which a finely tuned level of activity during S-phase is required to maintain fork speed, prevent replicative stress, and ensure the faithful segregation of the genetic blueprints during stem cell maintenance.

Given the prevalence of aneuploidy in early embryos and in cancer, it would be critical to explore whether low dosage of GSK3 inhibitors can be safety deployed in i) the fertility clinic to prevent embryo lost during *in vitro* fertilisation and ii) chromosomally unstable tumours for the prevention of their genomic evolution and chemotherapy resistance.

## Author Contributions

Sergio P. Acebrón and Anchel de Jaime-Soguero designed the research. Anchel de Jaime-Soguero conceived, performed and analysed experiments. Anna Jauch performed the karyotype studies. Sabine Costagliola, Mirian Romitti and Gayathri Vilangappurath performed experiments using hESCs (H1 and H9). Karl Willert contributed reagents and shared unpublished data. Floris Foijer assisted with the sequencing experiments. Sergio P. Acebrón supervised the project and wrote the paper with input of the other authors.

## Acknowledgements

We thank L. Villacorta and V. Benes (EMBL Heidelberg) for their assistance in sequencing experiments. We thank J. Hattemer and A. Ciprianidis for reagents and technical support. This work was supported by the Deutsche Forschungsgemeinschaft (DFG) SFB 1324, project number 331351713 (Project B03 to S.P.A.). J.H. was supported by a Studienstiftung PhD fellowship. A.d.J.S. was supported by an Actions Blanches 2024 grant (ULB) and the Mandat d’Impulsion Scientifique Ulysse (MISU) from the FNRS (project number MISU F.6002.26).

## Methods

### Cell culture

The hiPSCs were a gift from Kyung-Min Noh (EMBL). Cells were seeded in wells coated with Vitronectin 1 h at 37 °C (VTN 1:100 diluted in PBS), and grew in hiPSC culture medium. In detail, cells were cultured in Essential E8 medium (Thermo Scientific) supplemented with Penicillin/Streptomycin. We added Revitacell Supplement (Thermo Scientific) for the first 24 h after plating. Media were changed every day, and cells were split every 3–4 days using Versene (Gibco). All the experiments were initiated with hiPSCs between passages 10-13. For immunofluorescence experiments, 50.000 cells were seeded per each 12-well.

H9 and H1 hESCs experiments were conducted at IRIBHM-ULB in the laboratory of Dr. Sabine Costagliola, who holds an MTA agreement for their use. Cells were cultured in 6-well plates pre-coated with 1.6% growth factor reduced matrigel (356231, Corning) diluted in DMEM F12 (21331-020, Thermo Fisher Scientific) for 30 min at 37 °C in StemFlex medium (A3349401, ThermoFisher). hESCs were routinely passaged with EDTA 0.5 mM at a splitting ratio of 1–5 every 3-4 days, with media change every day.

For long term passage, hiPSC and hESC lines were routinely split during more than 90 days (30 passages) as explained above. Long term treatments included diverse GSK3 inhibitor CHIR99021 concentrations (500nM to 10nM range) and 20 μM nucleoside mix (dNs) (SCBT).

### Chromosome segregation analyses, ultrafine bridge quantification and unreplicated DNA in mitosis detection

Culture cells were fixed in 2–4% PFA for 10–15 min, permeabilised with 0.5% Triton X100 in PBS (PBST) for 10 min, followed by a blocking step for 20 min and overnight incubation with primary antibodies in 2% horse serum in 0.1% PBST. We used 1:250 guinea pig anti-CENPC antibody (MBL International Corporation, USA, cat no PD030) or CREST antibody to stain kinetochores (Chromosome segregation), or 1:250 mouse anti-phospho-H2AX (Ser139) (γ-H2AX) antibody (Millipore, clone JBW301) to detect DSBs, 1:250 mouse anti-phospho-RPA32/RPA2 (Ser4 + 8) (pRPA) antibody to detect single strand DNA breaks, 1:250 mouse anti-BLM (SantaCruz Biotech) to detect ultrafine bridges; and the secondary antibodies 1:500 anti-guinea pig Cy3 (Millipore, AP308P), 1:500 anti-mouse Alexa488 (ThermoFisher A21202), supplemented with 1 μg/ml DAPI. In all the figures where DNA damage is measured (gH2AX, pRPA), incorporated EdU was subjected to a click-it reaction (ThermoFisher, C10337), as indicated by the manufacturer.

H9 hESCs were platted at a ratio of 1:15 in 12 well plates on the top of 15 mm coverslips pre-sterilized in ethanol 100%, washed in sterile ddH2O and coated with 1.6% Growth Factor Reduced Matrigel diluted in DMEM F12. Two days after plating, cells were treated with 250 ng/mL of DKK1; 40 ng/mL of FGF2, or 200 ng/mL of Noggin for 16 h. Cells were fixed using 4% paraformaldehyde (PFA) (15710, Electron Microscopy Sciences) diluted in PBS for 20 min at room temperature and then washed three times with PBS—0.1% Tween. The permeabilisation step was performed in PBS containing 0.3% Triton X-100 and 0.1 M glycine for 30 min at room temperature. Samples were incubated with primary antibodies (CENPC— MBL and alternatively anti-Centromere Protein Antibody (CREST, 15-234 Antibodies Inc) diluted at 1:250 in blocking solution (3% BSA and 0.1% Tween in PBS) overnight at 4 °C. The day after, samples were washed with PBS—0.1% Tween three times, followed by incubation with secondary antibodies (donkey anti-guinea pig 594, AP193SA6 Millipore, and Goat anti-Human IgG (H+L) Cross-Adsorbed Secondary Antibody, Alexa Fluor™ 594 donkey anti-mouse 488, A-11014 ThermoFisher Scientific), diluted at 1:500 in blocking solution for 2 h at room temperature.

To quantify cells exhibiting chromosome missegregation, we analysed 70-100 anaphases (hiPSCs, hESCs) in each biological replicate using a Nikon Eclipse Ti using a 60× objective with oil immersion; or an inverted SP8 confocal microscope (Leica Microsystems) with a Leica 63×/1.4NA Oil objective. In hESCs and hiPSCs and their derived lineages, chromosomes clearly separated (Lagging) from the bulk of segregated DNA chromatids were considered as chromosome missegregation, both centric and acentric based on centromeric chromosome stainning. Basal levels of chromosome missegregation in hiPSCs and hESCs were in accordance with previous estimates [19, 24, 25].

To quantify the percentage of ultrafine bridges in hiPSC anaphases, we used BLM (Bloom) antibody staining as follows: cells were fixed with 2 % (w/v) PFA at RT for 5 min, followed by treatment with ice-cold methanol for another 5 min at −20 °C. Subsequently the slides were washed once with PBS and blocked by adding blocking solution (5 % (v/v) HS in PBS) for 30 min at RT. BLM was detected by using BLM antibody (SCBT, 1:500 in 2 % (v/v) FCS in PBS). Cells were incubated for 1.5 hours at RT. After 3 washing steps with PBS, cells were incubated with appropriate secondary antibodies conjugated to Alexa-Fluor488 (1:1000 in 2 % (v/v) HS in PBS) for 1 hour at RT. Subsequently, DNA was stained with Hoechst33342 (1:20,000 in PBS) for 5 min at RT. Finally, cells were washed three times with PBS, dried and mounted onto glass slides with VectaShield (Vector Laboratories, Burlingame, USA). At least 50 anaphase cells were determined per sample in three or more biological replicates.

To characterise the presence of unreplicated DNA in G2-early mitosis, we pulsed hiPSCs with EdU for 15 minutes. After fixation, we stained cells for mitotic marker phosphor-histone 3 (pH3) and perform the EdU Click-iT (C10337, ThermoFischer) reaction following manufacturer protocols.

### Primitive streak induction from hPSCs

Primitive streak-like cells were generated modifying previously described protocols [24, 34]. Briefly, we plated hiPSCs in low confluency (1:25; around 50.000 cells per 12 well) for 2 days in a hiPSC culture medium prior to differentiation. On the third day, we change the medium to A-RPMI media supplemented with GlutaMax and Penicillin/Streptomycin (Gibco) supplemented with 5 μM CHIR99021 and low doses of Activin-A (20ng/mL) for 24h and harvested the cells.

### Karyotype M-FISH analyses

Cells were treated for 16 h, as indicated, followed by 16 h arrest using 100 ng/mL nocodazole. M-FISH was performed as previously described [24]. Briefly, seven pools of flow-sorted human whole chromosome painting probes were amplified and combinatorial labelled using DEAC-, FITC-, Cy3, TexasRed, and Cy5-conjugated nucleotides and biotin-dUTP and digoxigenin-dUTP, respectively, by degenerative oligonucleotide-primed (DOP) PCR. Prior to hybridisation, metaphase spreads fixed on glass slides were digested with pepsin (0.5 mg/mL; Sigma) in 0.2 N HCL (Roth) for 10 min at 37 °C, washed in PBS, post-fixed in 1% formaldehyde, dehydrated with a degraded ethanol series and air dried. Slides were denatured in 70% formamide/1x SSC for 2 min at 72 °C. Hybridization mixture containing combinatorial labelled chromosome painting probes, an excess of unlabelled cot1 DNA in 50% formamide, 2× SSC, and 15% dextran sulphate were denatured for 7 min at 75 °C, pre-annealed for 20 min at 37 °C, and hybridized to the denatured metaphase preparations. After 48 h incubation at 37 °C slides were washed three times at room temperature in 2× SSC for 5 min, followed in 0.2× SSC/0.2% Tween-20 at 56 °C two times for 7 min. For indirect labelled probes, an immunofluorescence detection was carried out. Therefore, biotinylated probes were visualized using three layers of antibodies: streptavidin Alexa Fluor 750 conjugate (Invitrogen), biotinylated goat anti avidin (Vector) followed by a second streptavidin Alexa Fluor 750 conjugate (Invitrogen). Digoxigenin-labelled probes were visualized using two layers of antibodies: rabbit anti-digoxin (Sigma) followed by goat anti-rabbit IgG Cy5.5 (Linaris). Slides were washed three times in 4× SSC/0.2% Tween-20 for 5 min, counterstained with 4.6-diamidino-2-phenylindole (DAPI) and covered with anti-fade solution. Images of metaphase spreads were taken for each fluorochrome using highly specific filter sets (Chroma Technology, Brattleboro, VT) on a DM RXA epifluorescence microscope (Leica Microsystems, Bensheim, Germany) equipped with a Sensys CCD camera (Photometrics, Tucson, AZ). Camera and microscope were controlled by the Leica Q-FISH software and images were processed on the basis of the Leica MCK software and presented as multicolour karyograms (Leica Microsystems Imaging solutions).

### qRT-PCR

RNA was extracted and purified using a Quiagen RNAeasy Plus column kit, according to the manufacturer’s instructions. The cDNA was produced with SensiFAST cDNA Synthesis kit (Bioline) starting from 300 ng to 1 μg mRNA. Real-time quantitative PCR reactions from 8.3 ng of cDNA were set up in technical triplicate using the SensiFAST SYBR Hi-ROX kit (Bioline) on a StepOne Plus (ThermoScientific) and/or qTower qPCR machine. The sequences of the oligonucleotides used in this study are provided on request. Expression levels were normalized to PCR amplification with primers for beta Tubulin.

### RNA-sequencing preparation and analysis

hiPSCs cells were seeded in 6-well plates and cultured during 1 passage (hiPSCs p.0) or for 30 passages untreated, treated with GSK3i 100nM or nucleosides 20uM. As a positive control for WNT activation and pluripotency exit, hiPSCs were differentiated towards primitive streak-like cells at earky passages (*N* > 3 biological independent experiments per condition). RNA was extracted by using the RNeasy Mini Kit (QIAGEN). RNA integrity (RIN value) of all samples was assessed in an Agilent 2100 Bioanalyzer machine using an Agilent RNA 6000 Nano chip. After, library preparation of the RNA samples was performed using a NEBNext Ultra II RNA Library Prep Kit for Illumina. cDNA library quality and concentration were measured in an Agilent 2100 Bioanalyzer machine using an Agilent High Sensitivity DNA chip. After, libraries from all barcoded samples were pooled and sequenced. Sequencing reads were quality controlled using FastQC (http://www.bioinformatics.babraham.ac.uk/projects/fastqc/) tool (version 0.11.5). Read alignments, as well as count tables of mapped read per gene, were obtained by using STAR version 2.6.0a (Dobin *et al*, 2013) with GRCh38 human reference genome an its gene model (GRCh38.84). The dataset remains to be uploaded in a repository.

Differentially expressed genes were identified with DESeq2 version 2_1.20 (Anders & Huber, 2010) in R (See Dataset EV1A,B). The significant up- and downregulated genes (FDR < 0.1) were plotted in Volcano plots and heatmaps, and used for pathway and gene ontology (GO) (The Gene Ontology Consortium, 2019) enrichment analysis.

---

**Figure S1:**
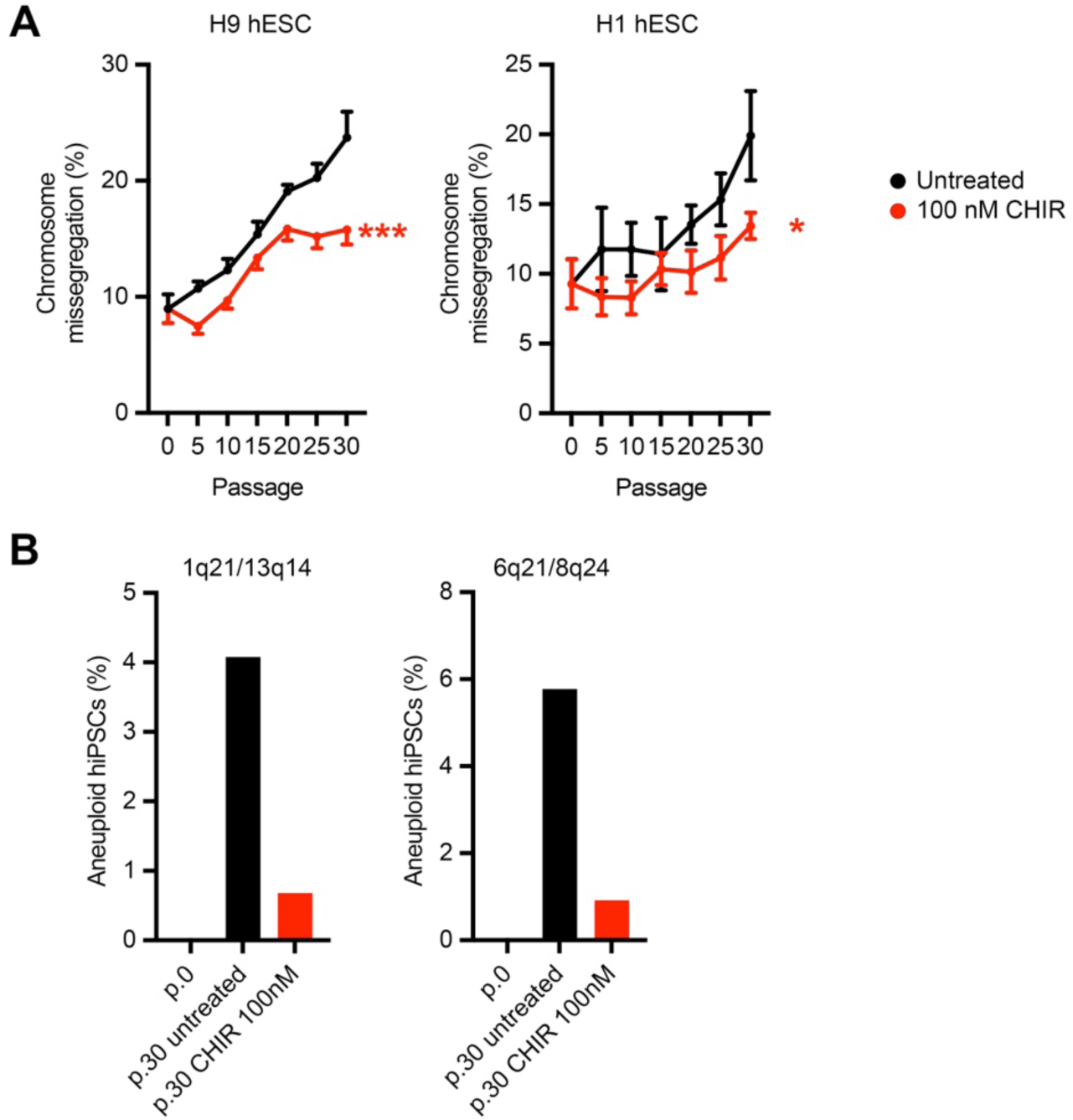
Chemical regulation of GSK3 activity prevents mitotic aberrations and aneuploidy in hPSCs. **A**, Chromosome segregation studies in hESCs expanded in E8 media in the presence or absence of 100 nM CHIR99021 (CHIR), and analysed at the indicated passages. **B**, Interphase FISH studies of hiPSCs cultured as indicated. Note that 2 pairs of chromosomes are analysed in each set.

**Figure S2:**
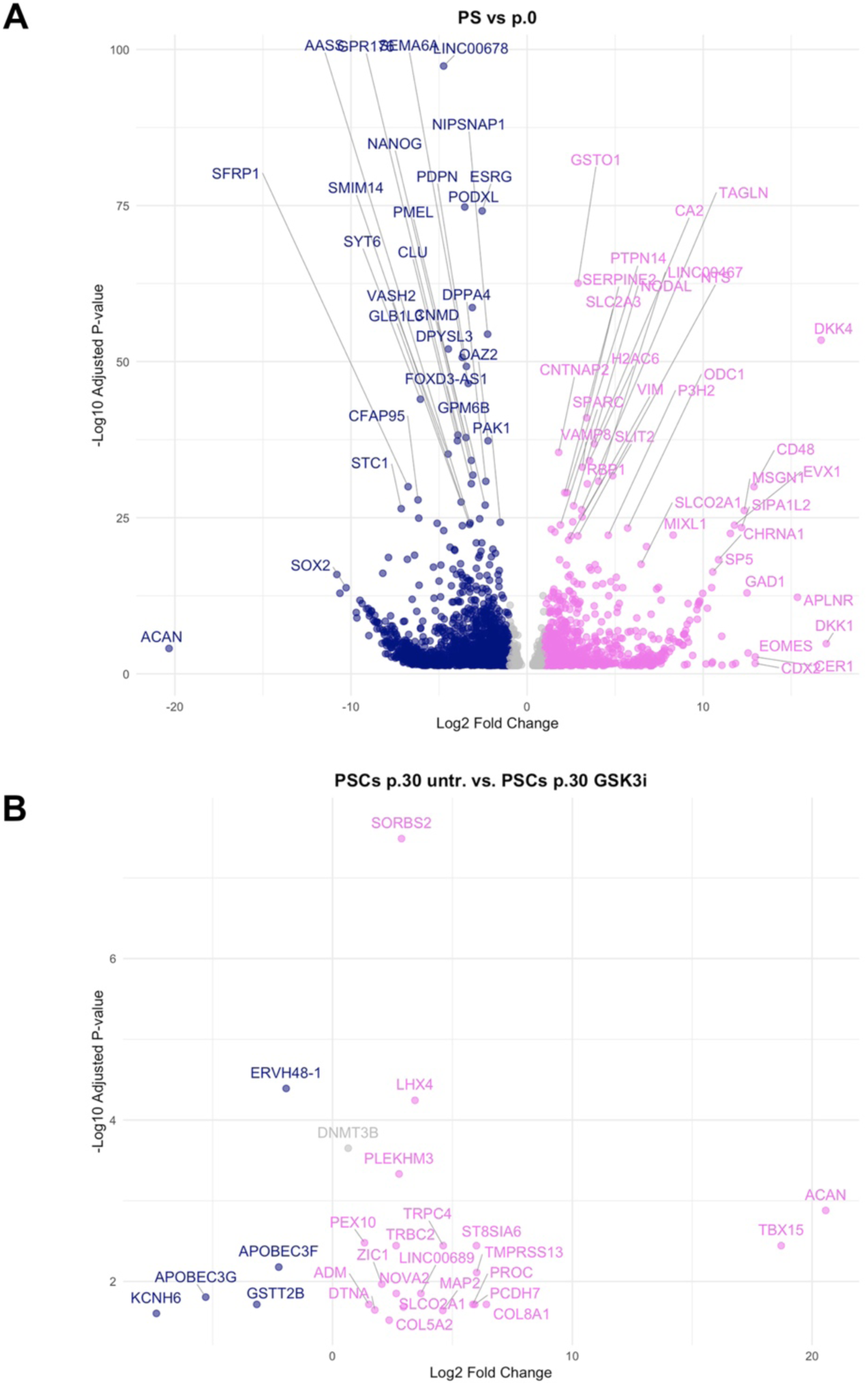
100 nM CHIR99021 does not impact hiPSC cell fate. **A**,**B**, Volcano plots of bulk sequencing analyses of human primitive-streak-like cells (PS), and hiPSCs cultured for up to 30 passages in E8 in the presence or absence of 100 nM CHIR99021 (GSK3i). Note that supplementation of GSK3i does not induce major transcriptional changes in targeted hiPSCs.

